# Female reproductive dormancy in *Drosophila melanogaster* is regulated by DH31-producing neurons projecting into the *corpus allatum*

**DOI:** 10.1101/2022.05.28.492955

**Authors:** Yoshitomo Kurogi, Eisuke Imura, Yosuke Mizuno, Ryo Hoshino, Marcela Nouzova, Shigeru Matsuyama, Akira Mizoguchi, Shu Kondo, Hiromu Tanimoto, Fernando G. Noriega, Ryusuke Niwa

**Affiliations:** Degree Programs in Life and Earth Sciences, Graduate School of Science and Technology, University of Tsukuba, Tennodai 1-1-1, Tsukuba, Ibaraki 305-8572, Japan; Graduate School of Life and Environmental Sciences, University of Tsukuba, Tennodai 1-1-1, Tsukuba, Ibaraki 305-8572, Japan; Life Science Center for Survival Dynamics, Tsukuba Advanced Research Alliance (TARA), University of Tsukuba, Tennodai 1-1-1, Tsukuba, Ibaraki 305-8577, Japan; Department of Biological Sciences and BSI, Florida International University, 11200 SW 8th street, Miami, FL 33199, United States of America; Biology Center of the Academy of Sciences of the Czech Republic, Institute of Parasitology, 37005, České Budějovice, Czech Republic; Faculty of Life and Environmental Sciences, University of Tsukuba, Tennodai 1-1-1, Tsukuba, Ibaraki 305-8572, Japan; Division of Liberal Arts and Sciences, Aichi Gakuin University, 12 Araike, Iwasaki-cho, Nisshin, Aichi 470-0195, Japan; Department of Biological Science and Technology, Faculty of Advanced Engineering, Tokyo University of Science, Niijuku 6-3-1, Katsushika-ku, Tokyo 125-8585, Japan; Invertebrate Genetics Laboratory, National Institute of Genetics, Yata 111, Mishima, Shizuoka 411-8540, Japan; Graduate School of Life Sciences, Tohoku University, Katahira 2-1-1, Sendai, Miyagi 980-8577, Japan; Department of Parasitology, University of South Bohemia, České Budějovice, Czech Republic

**Author notes:** Department of Entomology, Institute for Integrative Genome Biology, University of California, Riverside, 900 University Ave. Riverside, CA 92521, United States of America. Correspondence: Ryusuke Niwa, **Email:**. **Author Contributions:** Y.K., E.I., and R.N. designed the research; Y.K., E.I., Y.M., R.H., M.N., S.M., A.M., S.K., H.T., and F.G.N. performed the research; Y.K., E.I., M.N., and F.G.N. analyzed the data; Y.K. and R.N. wrote the paper; Y.K., E.I., H.T., F.G.N., and R.N. obtained the funding. **Competing Interest Statement:** The authors declare no competing or financial interests.

**Keywords:** reproductive dormancy, insect, juvenile hormone, corpus allatum, diuretic hormone 31

## Abstract

Female reproductive dormancy in insects is a process that drastically suppresses oogenesis to conserve energy under adverse environments. In many insects, including the fruit fly, *Drosophila melanogaster*, reproductive dormancy is induced under low-temperature and short-day conditions by the downregulation of juvenile hormone (JH) biosynthesis by the *corpus allatum* (CA). Previous studies have suggested that brain neurons that project directly to the CA are important for the regulation of reproductive dormancy. However, the role of CA-projecting neurons in JH-mediated reproductive dormancy has not yet been confirmed by molecular genetic studies. In this study, we report that, in adult *D. melanogaster*, the neuropeptide diuretic hormone 31 (DH31) is produced by brain neurons that project into the CA. DH31-producing-CA-projecting neurons are connected downstream with a subset of circadian clock neurons, such as s-LNvs, which are known to be involved in reproductive dormancy regulation. The CA expresses the gene encoding the DH31 receptor, which is required for DH31-triggered elevation of intracellular cAMP in the CA. Knocking down *Dh31* in these CA-projecting neurons or *DH31 receptor* in the CA leads to a failure in the decrease of the JH titer, normally observed under dormancy-inducing conditions, leading to abnormal yolk accumulation in the ovaries. Our findings provide the first molecular genetic evidence demonstrating that CA-projecting peptidergic neurons play an essential role in regulating reproductive dormancy by suppressing JH biosynthesis.

**Significance Statement:** Dormancy is an adaptive physiological response to environmental changes that are unsuitable for survival. Adult females of many insect species undergo reproductive dormancy in which oogenesis is drastically arrested; it is induced by a decrease in juvenile hormone (JH) titers. However, we are yet to fully understand the molecular mechanisms underlying the control of JH biosynthesis under dormancy-inducing conditions. In this study using the fruit fly, we demonstrated that brain neurons projecting directly to the JH-producing organ, *corpus allatum*, play an essential role in regulating reproductive dormancy via the neuropeptide DH31. As the morphologically-similar neurons have previously been suggested to be involved in reproductive dormancy regulation, this study provides a fundamental molecular and neuronal basis for reproductive dormancy in insects.

## Introduction

Unfavorable seasonal changes for prolonged durations, such as extremely low temperatures and food shortages during winter, may be challenging for the survival of animals in the temperate zone. During such unfavorable conditions, organisms often suspend or retard their normal development, growth, and physiological functions, known as dormancy (1). Dormancy has been intensively studied in insects as its control could benefit many aspects of industry and agriculture (2–4). Previous studies have revealed that a fraction of insects enter dormancy at the adult stage under dormancy-inducing conditions leading to multiple metabolic and behavioral changes, including slowed or halted reproductive maturation known as reproductive dormancy (4–6). As female insect adults allocate extensive energy resources toward oogenesis (7), reproductive dormancy allows them to reduce their energy consumption and resume reproduction under favorable conditions.

Reproductive dormancy in insects is regulated by a complex interplay between multiple hormones and neurotransmitters (3–5). Among these, reduction in the titer of juvenile hormone (JH), an insect-specific sesquiterpenoid hormones (8, 9), has been extensively studied, revealing its vital role in regulating reproductive dormancy in female adult insects (5). Since JHs are essential for promoting vitellogenesis under non-dormancy-inducing conditions, a reduction in hemolymph JH levels is required for suppressing vitellogenesis, leading to reproductive dormancy in females (3, 10). In many insect species, the reduction of hemolymph JH levels correlates with the downregulation of JH biosynthesis in a specialized endocrine organ called the *corpus allatum* (CA)(1, 3, 5). Previous studies have demonstrated that surgical amputation of the nervous connection between the brain and the CA impairs the induction of reproductive dormancy (11, 12). These observations suggest the presence of a mechanism of signal transduction by which information about unfavorable environmental conditions are processed in the brain and transmitted to the CA to reduce JH biosynthesis.

Previous studies have demonstrated that brain neurons projecting to the CA play crucial roles in the induction of reproductive dormancy in multiple insect species (3, 4, 13). For example, some neurons whose cell bodies are located in the anterior midline of the brain, known as *pars intercerebralis* (PI), project to the CA in the common ground-hopper, *Tetrix undulata* (14); linden bug, *Pyrrhocoris apterus* (11); blowfly, *Protophormia terraenovae* (15, 16); and brown-winged green bug, *Plautia stali* (17). A study reported that specific PI neurons in *P. stali* produce *P. stali* myoinhibitory peptide (Plast-MIP), a neuropeptide that inhibits JH biosynthesis in CA under dormancy-inducing conditions (18–20). It has also been demonstrated that a group of neurons, whose cell bodies are located on the lateral side of the brain, known as *pars lateralis* (PL), project to the CA in *P. terraenovae* (15); the bean bug, *Riptortus pedestris* (21); Colorado potato beetle, *Leptinotarsa decemlineata* (22); and grasshopper, *Locusta migratoria* (23). In all these cases, PL neurons seem to play an inhibitory role in ovarian development under dormancy-inducing conditions. Moreover, cauterization experiments in *L. decemlineata* and *L. migratoria* have suggested that PL plays an inhibitory role in JH biosynthesis under dormancy-inducing conditions (23, 24). In *P. terraenovae*, specific PL neurons projecting to the CA produce neuropeptides including cholesistokinin-8 and FMRF-amide (25, 26). However, despite their crucial role in regulating reproductive dormancy, the role of CA-projecting PL neurons in reproductive dormancy has not been genetically confirmed. Moreover, it is unclear which neurotransmitters or neuropeptides in CA-projecting PL neurons are responsible for regulating reproductive dormancy in insect species.

Studies on the fruit fly, *Drosophila melanogaster*, have contributed substantially to the discovery of several regulatory mechanisms of reproductive dormancy (5). Similar to other insects, reproductive dormancy in *D. melanogaster* is induced under low-temperature and short-day conditions (27). Several studies have confirmed that JH biosynthesis and signaling are essential for the regulation of reproductive dormancy in *D. melanogaster* (28–30). In this study, we report the identification and characterization of CA-projecting PL neurons that produce the neuropeptide diuretic hormone 31 (DH31). DH31 is known to play versatile roles in insects, particularly in *D. melanogaster*, such as diuretic action, establishment of daily temperature preference rhythms, locomotor activity, sleep regulation, midgut contraction frequency, intestinal immune responses, and nutrient-dependent regulation of courtship (31–40). In this study, we propose that DH31 signaling to CA plays an important role in the inhibition of JH biosynthesis under dormancy-inducing conditions, leading to the induction of reproductive dormancy through the inhibition of JH-mediated maturation of eggs.

## Results

### A subset of DH31-producing neurons projects into the *D. melanogaster corpus allatum*

In our previous studies, we have identified several neurons projecting to various endocrine organs, including the CA of *D. melanogaster* (41–43). To further identify CA-projecting neurons, we conducted immunostaining experiments using antibodies against some neuropeptides, and investigated whether any of these peptidergic-cell neuronal processes innervate the CA, as described previously for the hugin-positive neurons (43). Our results demonstrated that DH31-immunoreactive fibers and varicosities were present within the CA (Fig. 1A, B). A trace back of these CA-projecting DH31-producing neurons (Fig. 1A) led to three pairs of cell bodies located on the dorsal side of the central brain (Fig. 1A, C). Among these, two cell bodies in the brain hemisphere were clustered, whereas another cell body was located closer to the midline. We also found that the *R21C09-GAL4* driver, in which the *GAL4* transgene is expressed under the control of part of *Dh31* enhancer, labeled these CA-projecting neurons. The *R21C09-GAL4*-driven GFP signals were observed in three pairs of the central brain neurons and neuronal processes in the CA region, which were co-immunostained with anti-DH31 antibody (Fig. S1A, B). Hereafter, we designate these three pairs of CA-projecting DH31-producing neurons as “DH31_CA_ neurons.”

**Figure 1.**
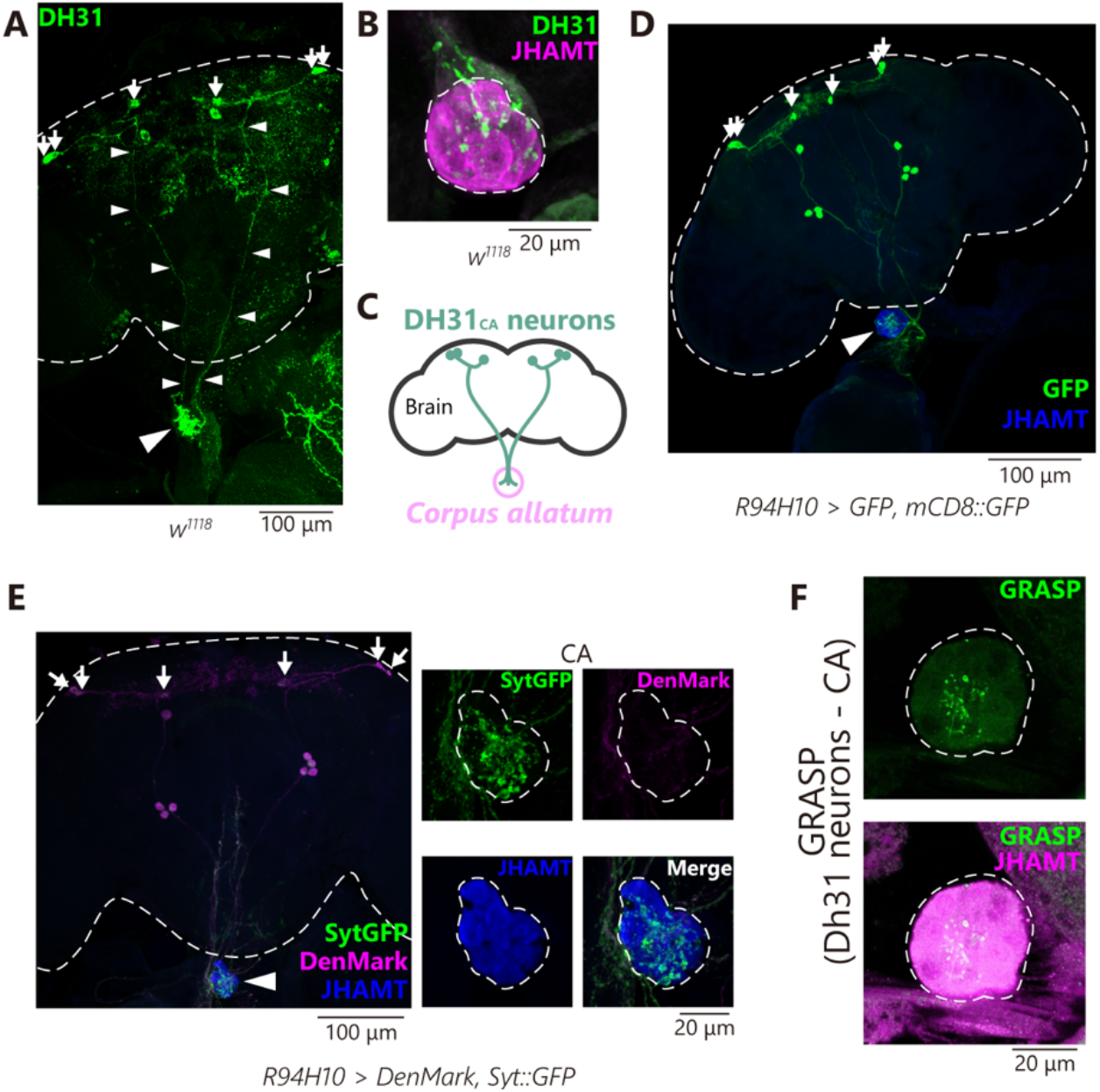
Anatomical characterization of DH31_CA_ neurons; a subset of DH31-producing neurons project to the CA. In all panels, small arrows indicate the cell bodies of DH31_CA_ neurons. (A) Immunostaining with anti-DH31 antibody in the adult central brain (outlined by dashed lines) and the CA of wild-type (*w*^*1118*^) adult females. The upper and lower parts of the photograph correspond to the dorsal and ventral sides of the central brain, respectively. Axonal processes from the cell bodies to the CA (large arrowhead) are indicated by the small arrowheads. (B) Immunostaining signal in the CA of wild-type (*w*^*1118*^) adult females four d after eclosion. The anti-DH31 antibody (green) was employed along with the anti-JHAMT antibody, which was used to visualize the CA (magenta). DH31-positive puncta are observed in the CA region. Note that DH31-immunoreacive varicosities were observed inside the CA. (C) A schematic drawing representing the anatomy of brain, DH31_CA_ neurons, and CA. (D) Transgenic visualization of DH31_CA_ neurons by GFP driven by *R94H10-GAL4*, which specifically labels DH31_CA_ neurons and other small subsets of neurons. Samples were immunostained with anti-GFP (green) and anti-JHAMT (blue) antibodies. The CA is marked with a large arrowhead. (E) (Left) Visualization of axons and dendrites of DH31_CA_ neurons stained by synaptotagmin-GFP (SytGFP, green) and DenMark (magenta) driven by *R94H10-GAL4*. The CA (large arrowhead) is visualized by immunostaining with anti-JHAMT (blue). (Right, four small panels) magnified view of the left panel focusing on the CA region. SytGFP, but not DenMark, signals were observed in the CA region, indicating that axons of DH31_CA_ neurons innervate the CA. (F) The GRASP signal was employed to visualize the close physical contact between DH31_CA_ neurons and the CA. Also see Fig. S1G for the GRASP negative controls.

A previous study identified one pair of CA-projecting lateral protocerebrum neurons #1 (CA-LP1 neurons) and two pairs of CA-LP2 neurons, which innervate the CA during the *D. melanogaster* larval stage (44, 45). However, the neurotransmitters produced by these neurons remain unknown. As the positions of the DH31_CA_ cell bodies were similar to those of CA-LP1 and CA-LP2 neurons, we examined whether the CA-LP1 and CA-LP2 neurons were identical to the DH31_CA_ neurons. We found that *Kurs21-GAL4*-driven GFP, which reportedly marks CA-LP1 and CA-LP2 neurons (44) Fig. S1C), visualized the projection of their neurites to the CA, in the wandering third-instar larvae (Fig. S1D) as well as in adults (Fig. S1E, F). Moreover, the cell bodies and CA-projecting neurites of CA-LP1 and CA-LP2 neurons were immunostained with the anti-DH31 antibody in both larval and adult stages (Fig. S1C–F). Therefore, DH31_CA_ neurons in the *D. melanogaster* adult stage are identical to the CA-LP1 and CA-LP2 neurons described in its larval stage.

*R21C09-GAL4* and *Kurs21-GAL4* were active in many neurons, apart from DH31_CA_ neurons, in the central brain (Fig. S1G, H). We thus searched for a *GAL4* driver that would be active in a restricted group of neuronal cells including the DH31_CA_ neurons. We browsed large collections of adult fly images from the FlyLight database of the Janelia Research Campus, Howard Hugh Medical Institute (https://flweb.janelia.org/cgi-bin/flew.cgi). The FlyLight database provides large anatomical image datasets and a well-characterized collection of *GAL4* lines, allowing us to visualize individual neurons in the *D. melanogaster* central nervous system (46). We manually checked most of the images of *GMR GAL4* lines deposited in FlyLight and found *R94H10-GAL4* labeled DH31_CA_ neurons and a few additional neurons (Fig. 1D, Fig. S2A, B). In addition to DH31_CA_ neurons, other *R94H10-GAL4*-positive neurons were DH31-negative (Fig. S2A). Furthermore, while *Dh31* is expressed in enteroendocrine cells (47), *R94H10-GAL4* was not active in these gut cells (Fig. S2C). Therefore, in this study, we used the *GAL4* driver to manipulate gene expression in DH31_CA_ neurons.

Thereafter, we examined the distribution of axonal termini and dendrites in the DH31_CA_ neurons. *Synaptotagmin::GFP* (*Syt::GFP*) and *DenMark* transgenes, whose translated products were localized at the axonal termini and dendrites, respectively (48, 49), were driven by *R94H10-GAL4*. We found that the Syt::GFP and DenMark signals were primarily observed in the CA region and in the dorsal side of the central brain, respectively (Fig. 1E), suggesting that the DH31-immunoreactive fibers in the CA regions are axons. Furthermore, GFP reconstitution across synaptic partners (GRASP) analysis (50), in which two complementary fragments of GFP were expressed in DH31_CA_ neurons and the CA, respectively, revealed that GRASP signals were detected inside the CA (Fig. 1F, Fig. S3A, B). These results suggest that DH31_CA_ neurons project to the CA and that they are in physical contact with the CA.

### A subset of circadian clock neurons innervates the DH31_CA_ neurons

To anatomically characterize DH31 neurons, we browsed the publicly available connectome database of adult *D. melanogaster* brains to search for neurons that connect upstream to DH31_CA_ neurons (see Materials and Methods). In the available connectome data of neurons labeled with *R21C09-GAL4*, the two pairs of clustered DH31_CA_ neurons have been annotated (Fig. S4A–D). The anatomical positions of the cell bodies and axons of these neurons suggested that they correspond to the CA-LP2 neurons (44) Fig. S1C, D). In contrast, another pair of DH31_CA_ neurons that was similar to CA-LP1 neurons was not clearly identified in the available database. Therefore, in our connectome analysis, we focused on CA-LP2 neurons that exhibited 1780 connections with other neurons. Among these neurons, we found that several clock neurons, including the 5^th^ sLN-v, DN1p, LNd, and s-LNv, were connected to the CA-LP2 neurons (Fig. 2A). These results suggest that the circadian clock may modulate the function of the DH31_CA_ neurons.

**Figure 2.**
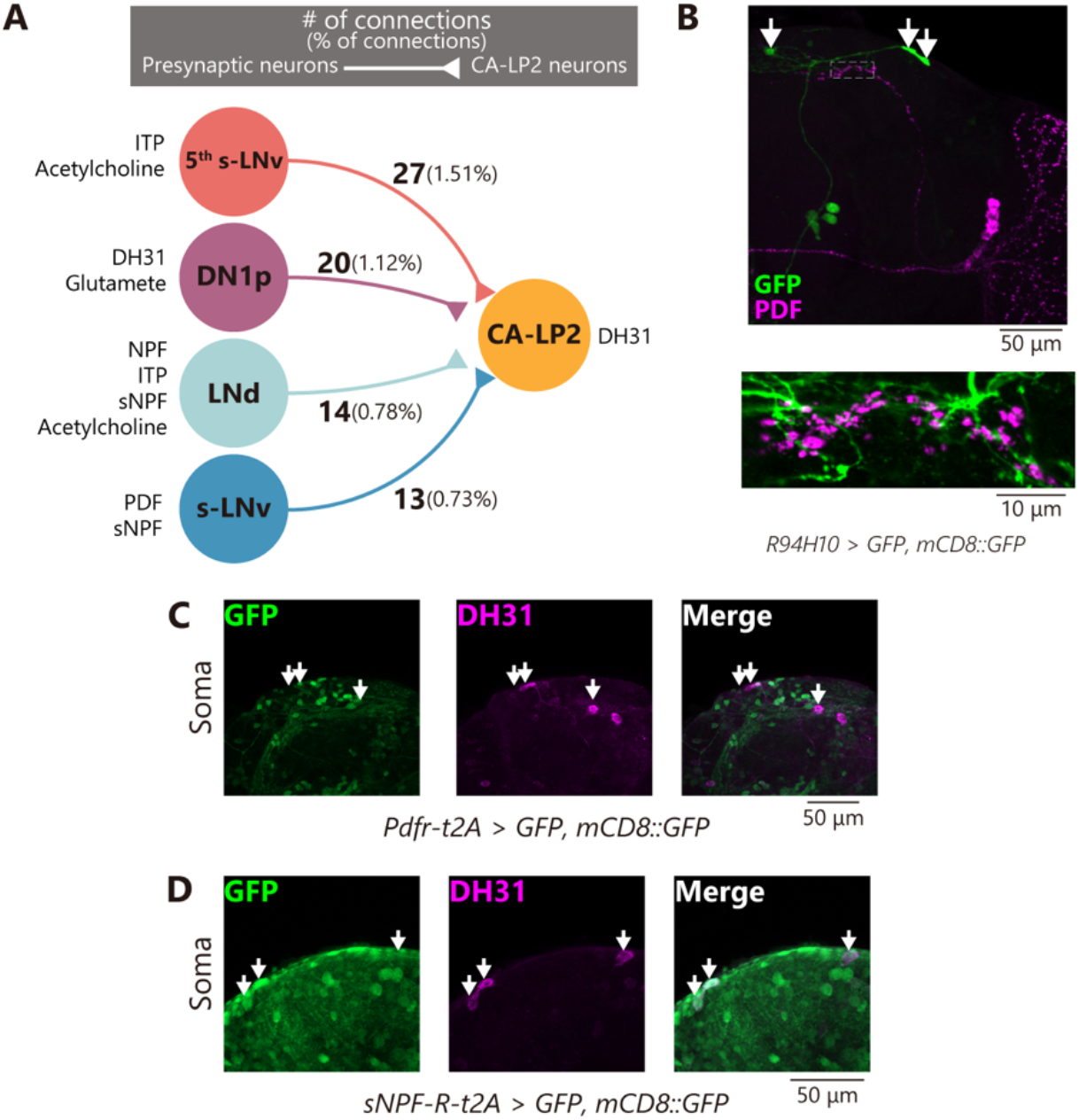
A subset of circadian clock neurons innervates the DH31_CA_ neurons. In all panels, arrows indicate the cell bodies of DH31_CA_ neurons. (A) Connectivity diagram between DH31_CA_ and clock neurons (s-LNv, 5^th^ s-LNv, DN1p, and LNd). Numbers outside parentheses indicate contact site counts marked by presynaptic densities. Numbers in parentheses indicate the percentage of contact site counts with each clock neuron population per total contact between DH31_CA_ neurons and all the neurons described in neuPrint+. Neuropeptides and neurotransmitters produced in the clock neurons are also indicated. (B) Immunostaining with anti-GFP (green) and anti-PDF (magenta) antibodies in an adult female expressing GFP driven by *R94H10-GAL4*. The lower panel represents high magnification, with increased gains in the dashed inset in the upper panel. Note that the axonal termini of PDF clock neurons are in close proximity to the neuronal processes of DH31_CA_ neurons. (C, D) Expression of *Pdfr* (C) and *sNPF-R* (D) visualized using *Pdfr-T2A-GAL4*-and *sNFPR-T2A-GAL4*-driven GFP, respectively (green). The soma of the DH31_CA_ neurons is marked with an anti-DH31 antibody (magenta).

s-LNvs are involved in regulating reproductive dormancy in *D. melanogaster* (51). Moreover, the s-LNvs innervate neurons located in the PL, which closely corresponds to the LP the region in the brain where the CA projecting neurons are located in several insects, including the blowfly, *P. terraenovae* (52–54). Therefore, we examined the relationship between s-LNvs and DH31_CA_ neurons in *D. melanogaster*. The morphology of s-LNvs can be visualized using the anti-pigment dispersing factor (PDF) antibodies (55). We found that the axonal termini of s-LNvs were similar to the neuronal processes of DH31_CA_ neurons (Fig. 2B, Fig. S4E). We initially expected that the PDF receptor (Pdrf) would be present in DH31_CA_ neurons; however, *Pdfr knock-in T2A GAL4* was absent in these neurons (Fig. 2C). In contrast, the short neuropeptide F (sNPF), which is known to be present in s-LNvs (56), might be involved, as the *s-NPF receptor* (*s-NPF-R*) *knock-in T2A GAL4* was positive in these neurons (Fig. 2D). These results suggest that DH31_CA_ neurons, to an extent, have chemical connections with s-LNvs, at least through sNPF.

### The DH31_CA_ neurons are required for inducing reproductive dormancy

Considering that DH31_CA_ neurons are located in the lateral protocerebrum and in proximity to the axon termini of s-LNvs, the morphological and anatomical features of *D. melanogaster* DH31_CA_ neurons are substantially similar to those of *P. terraenovae* CA-projecting neurons (16, 53). A previous study reported that CA-projecting neurons negatively regulate the induction of reproductive dormancy in *P. terraenovae*, as surgical amputation of the axons of CA-projecting neurons caused abnormal egg production, even under dormancy-inducing conditions (57). Therefore, we examined whether DH31_CA_ neurons were also involved in reproductive dormancy in *D. melanogaster*.

We first examined whether genetic mutants with a loss of *Dh31* function displayed any phenotypes of reproductive dormancy in virgin females. We confirmed that DH31 immunoreactivity in the CA region was diminished in the loss of *Dh31* function flies (Fig. 3A), suggesting that DH31 is not active in the CA of mutant insects. Furthermore, under dormancy-inducing conditions (at 11 ± 0.5 °C under 10:14 h light/dark cycle), the loss of *Dh31* function in females led to significant enlargement of the ovaries compared to those in control females that exhibited the typical dormancy-induced reduction of ovarian development (Fig. 3B). Finally, the loss of *Dh31* function in females led to mature egg production compared to the control females (Fig. 3C).

**Figure 3.**
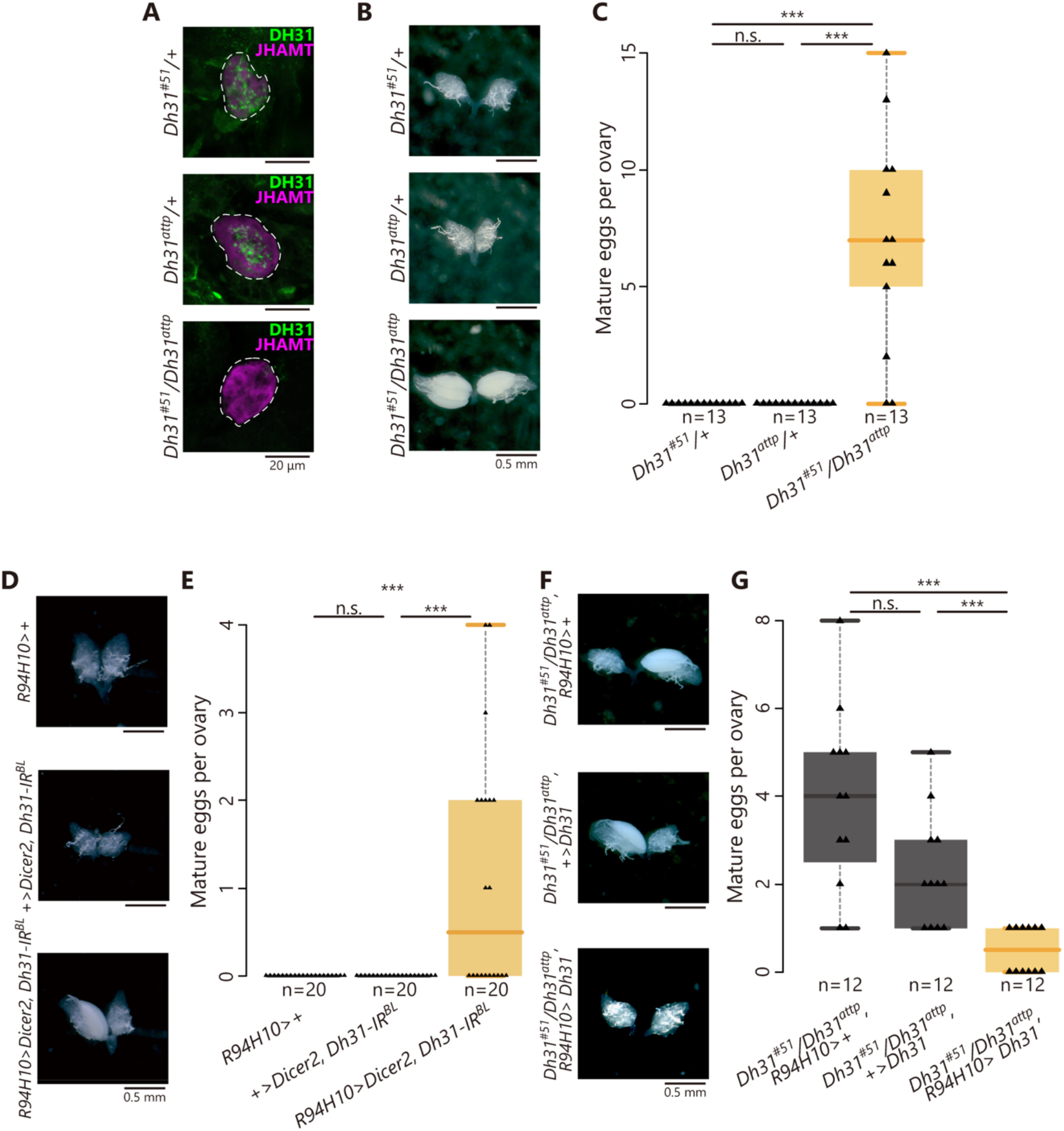
The loss of DH31 function in DH31_CA_ neurons fails to induce reproductive dormancy. Samples were derived from virgin females 12 d after their transfer to dormancy-inducing conditions. (A) Immunostaining signals of anti-DH31 antibody (green) in the CA region of control and *Dh31* genetic mutant females. CA was visualized with an anti-JHAMT antibody (magenta). Note that DH31 puncta were diminished in the CA region of *Dh31* trans-heterozygous mutant females. (B, C) Mature egg formation in control and *Dh31* genetic mutant females under dormancy-inducing conditions. (B) Representative images of the ovaries. (C) Quantification of mature eggs per ovary. (D, E) Mature egg formation in control and DH31_CA_ neuron-specific *Dh31* RNAi females under dormancy-inducing conditions. (D) Representative images of the ovaries. (E) Quantification of mature eggs per ovary. (F, G) Mature egg formation in *Dh31* trans-heterozygous mutant females with or without DH31_CA_ neuron-specific expression under dormancy-inducing conditions. (F) Representative images of the ovaries. (G) Quantification of mature eggs per ovary. Statistical analysis: Wilcoxon rank sum test with Bonferroni’s correction for C, E, and G. \*\*\**P* <0.001. n.s.: not significant.

We next examined whether the DH31 peptide, produced in DH31_CA_ neurons, was a critical regulator of reproductive dormancy. For this purpose, we knocked down *Dh31* specifically in DH31_CA_ neurons using a transgenic RNAi technique. An *R94H10-GAL4*-driven *Dh31* inverted repeat (IR) construct eliminated almost all the DH31 protein in the CA (Fig. S5A). Under dormancy-inducing conditions, RNAi-treated females displayed greater ovarian development and higher production of mature eggs than the control females (Fig. 3D, E). We implemented an additional genetic approach, in which a *Dh31* transgene was driven specifically in DH31_CA_ neurons in the genetic background of the loss of *Dh31* function. An *R94H10-GAL4*-driven *Dh31* overexpression restored DH31 protein levels in the CA of the mutant females (Fig. S5B). While the loss of *Dh31* function in females failed to induce reproductive dormancy, *Dh31* overexpression rescued the phenotypes of reproductive dormancy (Fig. 3F, G). Therefore, these results suggest that DH31 produced by DH31_CA_ neurons plays an essential role in the suppression of oogenesis under dormancy-inducing conditions.

### The DH31 receptor in the *corpus allatum* is required for the induction of reproductive dormancy

As DH31_CA_ neurons project to and are in close contact with the CA, we examined whether a specific receptor for DH31 (DH31-R) (58) was present in the CA. For this purpose, we generated a knock-in strain by inserting the *T2A-GAL4* cassette into the *Dh31-R* locus (49). This transgenic line was used to confirm the *Dh31-R* expression in the CA (Fig. 4A). We then conducted phenotypic analyses of the genetic mutants that presented a loss of DH31-R function. Under dormancy-inducing conditions, we observed impaired ovarian reduction and suppression of mature egg formation (Fig. 4B, C). Similar reproductive phenotypes were also observed in females in which *Dh31-R* was knocked down in the CA with *Aug21-GAL4*, known to be active in the CA (44, 45) (Fig. 4D, E). *Aug21-GAL4*-driven RNAi phenotypes were observed using two independent *Dh31-R* IR constructs (Fig. S6A, B). These results confirm that DH31-R in the CA is required for the induction of reproductive dormancy.

**Figure 4.**
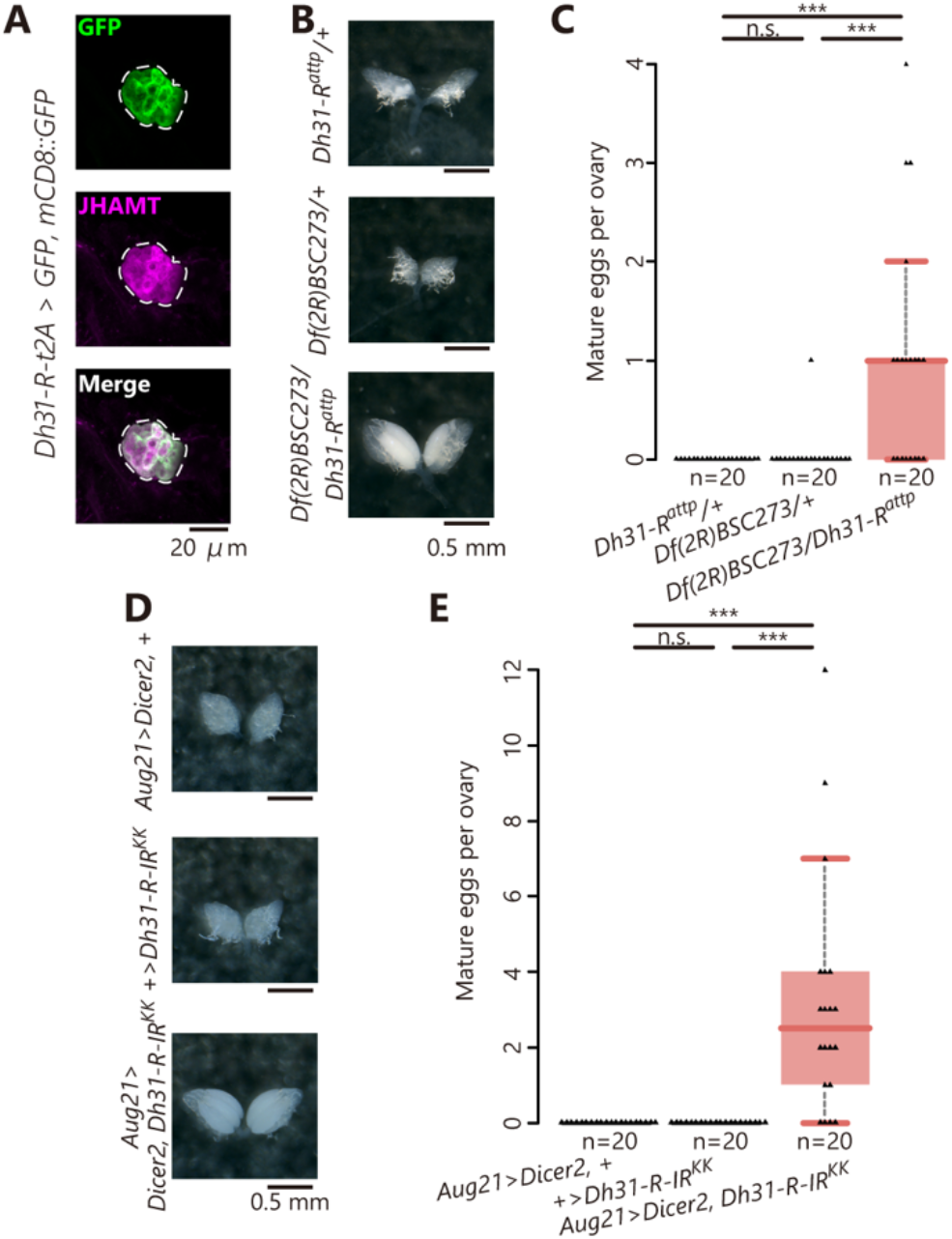
The loss of DH31 receptor function in the CA leads to failure in the induction of reproductive dormancy. Samples were derived from virgin females 12 d after their transfer to dormancy-inducing conditions. (A) Expression of *Dh31-R* visualized using *Dh31-R-T2A-GAL4*-driven GFP (green). CA was revealed using an anti-JHAMT antibody (magenta). (B, C) Mature egg formation in control and *Dh31-R* mutant females under dormancy-inducing conditions. (B) Representative images of the ovaries. (C) Quantification of mature eggs per ovary. (D, E) Mature egg formation in the control and CA-specific *Dh31-R* RNAi females under dormancy-inducing conditions. (D) Representative images of the ovaries. (E) Quantification of mature eggs per ovary. Note that these data were obtained from the Vienna Drosophila Resource Center KK RNAi strain. (Also see Fig. S4A and B, in which the results were obtained by employing a different transgenic RNAi construct.) Statistical analysis: Wilcoxon rank-sum test with Bonferroni correction for C and E. \*\*\**P* <0.001. n.s.: not significant.

### DH31 induces intracellular cAMP elevation in the CA through the DH31 receptor

We employed live imaging techniques to further understand the role of DH31-R in transmitting DH31 signals to CA cells. Considering that DH31-R is coupled to cAMP as a secondary messenger (58), we monitored the increase in intracellular cAMP levels using CA *ex vivo* culture system that assays the activity of the cAMP sensor probe, PinkFlamindo (59). Fly tissues containing the CA expressing the *PinkFlamindo* transgene were dissected and treated with collagenase because the CA was surrounded by a thick layer of collagens (Fig. S7A). The collagenase-treated tissue was placed in a glass-bottomed dish filled with culture medium (Fig. 5A, Fig. S7B); subsequently, the PinkFlamindo signals were recorded in the presence and absence of extracellular chemical stimuli. We first confirmed that the PinkFlamindo signals reflected intracellular cAMP levels, since the administration of the adenylyl cyclase activator, NKH477 elevated PinkFlamindo signals in the CA (Fig. S7C, D). Utilizing this *ex vivo* system, we found that the administration of the synthetic DH31 peptide resulted in the prompt elevation of PinkFlamindo signals in the CA (Fig. 5B). Importantly, the DH31-stimulated elevation of PinkFlamindo signals was not observed in the *Dh31-R* knocked down CA (Fig. 5D, E). These results indicate that DH31 signals via DH31-R, lead to an increase in intracellular cAMP levels in CA.

**Figure 5.**
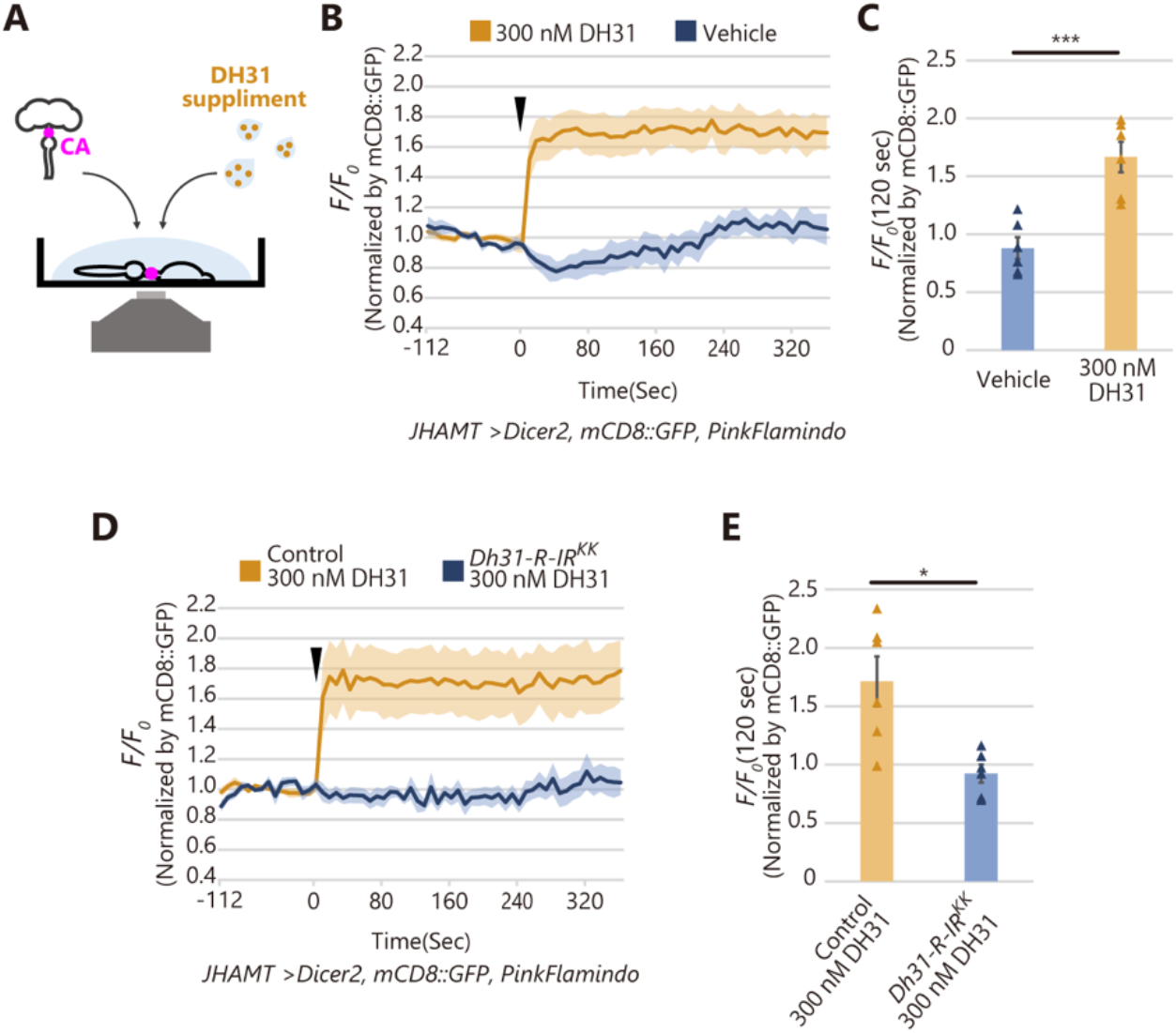
Synthetic DH31 peptide elevates intracellular cAMP level in the CA through the DH31 receptor. (A) A schematic representation of *ex vivo* cAMP imaging. The dissected tissues, containing the brain, gut, and CA, were incubated in Schneider’s *Drosophila* medium in the presence or absence of synthetic DH31 peptide. (B) Changes in the relative fluorescence intensity of *JHAMT-GAL4*-driven PinkFlamindo in the CA after 360 s with or without stimulation using 0.3 μM DH31 peptide (each n = 6). *F/F0* values are normalized by the signal intensity of *JHAMT-GAL4*-driven mCD8::GFP. (C) Quantification of normalized PinkFlamindo signals in the CA at 120 s after the stimulation. (D) Changes in the relative fluorescence intensity of *JHAMT-GAL4*-driven PinkFlamindo in the CA of control and *Dh31-R* RNAi females after 360 s with stimulation using 0.3 μM DH31 peptide (each n = 6). *F/F0* values are normalized by the signal intensity of *JHAMT-GAL4*-driven mCD8::GFP. (E) Quantification of normalized PinkFlamindo signals in the CA at 120 s after the stimulation. Values in B to E are presented as mean ± SE. Statistical analysis: Student’s *t*-test for C and E. \**P* <0.05, \*\*\**P* <0.001. n.s.: not significant.

### DH31 signaling in the CA plays a role in decreasing hemolymph JH titers

The induction of *D. melanogaster* reproductive dormancy is associated with the suppression of JH biosynthesis in the CA (29, 30, 60–62); therefore, we examined whether DH31 signaling in CA influenced JH titers in the hemolymph. We used liquid chromatography coupled with electrospray tandem mass spectrometry (LC-MS/MS) (63) to measure hemolymph titers of JH III, a major form of JH in *D. melanogaster* adult females (64–67). Females with the loss of *Dh31* or *Dh31-R* function exhibited a higher hemolymph JH III titer than the control females (Fig. 6A, B). In addition, under dormancy-inducing conditions, administration of the JH analog, methoprene induced egg maturation, which phenocopied females with loss of *Dh31* function (Fig. S8A, B).

**Figure 6.**
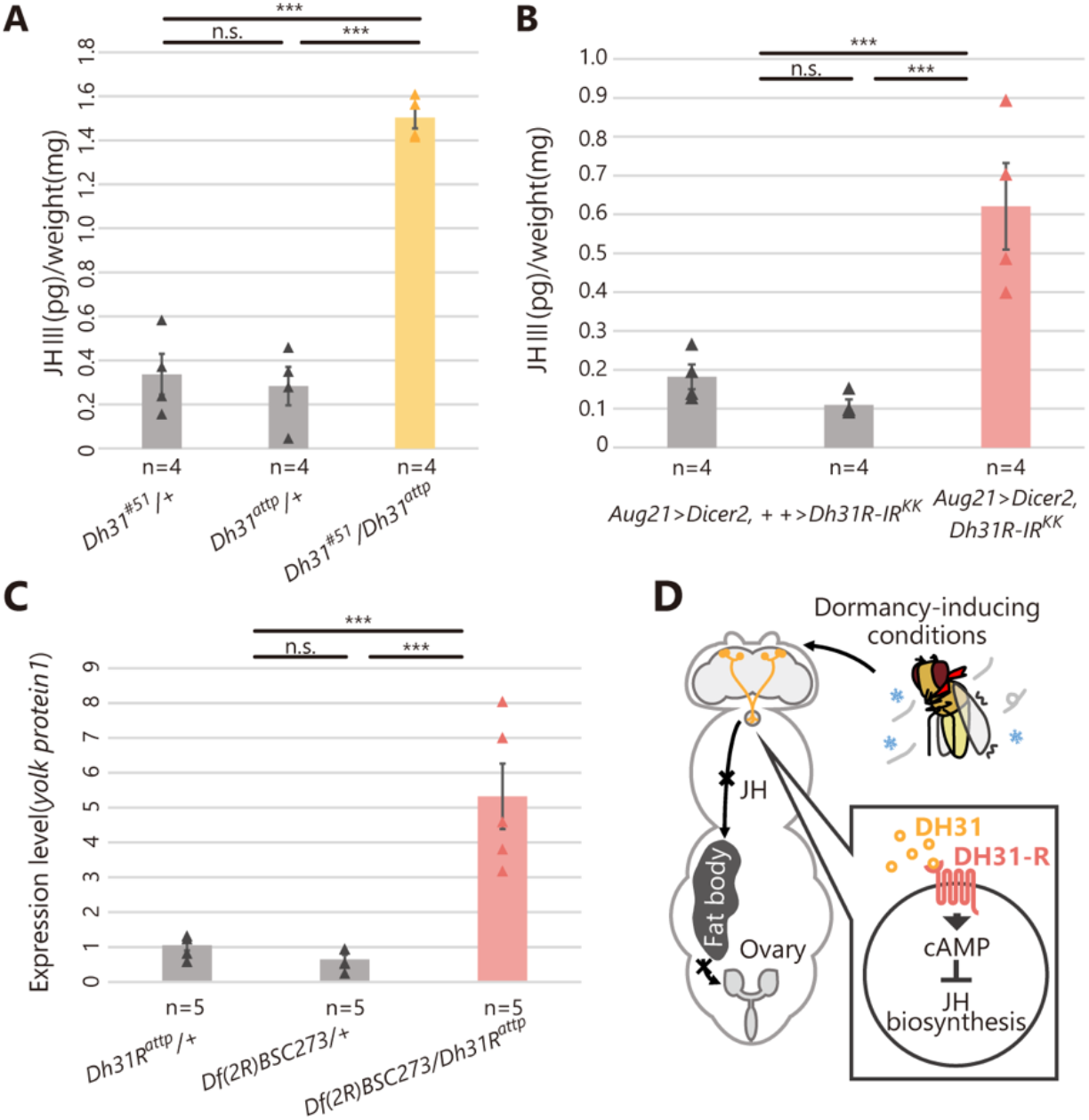
Hemolymph JH III titers are increased after the loss of DH31 signaling. Samples were derived from virgin females 6 d after their transfer to dormancy-inducing conditions. (A) Hemolymph JH III titers in control and *Dh31* genetic mutant females under dormancy-inducing conditions. JH III amounts are normalized by body weight. (B) Hemolymph JH III titers in control and CA-specific *Dh31-R* RNAi females under dormancy inducing conditions. JH III amounts are normalized by body weight. (C) Quantification of *yolk protein 1* mRNA in control and *Dh31-R* genetic mutant females under dormancy inducing conditions by RT-qPCR. (D) A proposed model of DH31-dependent regulation of reproductive dormancy in *D. melanogaster*. Values in A to C are presented as mean ± SE. Statistical analysis: Tukey–Kramer’s HSD test for G and H. *** *P* <0.001. n.s.: not significant.

JH promotes yolk protein synthesis in the fat body to promote mature egg production (68–70). Consistently, the mRNA levels of *yolk protein 1, yolk protein 2*, and *yolk protein 3* were upregulated in females with loss of *Dh31-R* function (Fig. 6C, Fig. S8C, D). Altogether, these results suggest that DH31 signaling in CA plays a crucial role in suppressing the hemolymph JH titer under dormancy-inducing conditions, leading to reproductive dormancy through the inhibition of JH-mediated oocyte maturation (Fig. 6D).

## Discussion

CA-projecting PL neurons are essential regulators of reproductive dormancy in insects (15, 21– 23). In addition, CA-projecting PL neurons are known to play a critical inhibitory role in JH biosynthesis (23, 24). The current study provides the first genetic evidence of the role of CA-projecting PL neurons in JH-mediated reproductive dormancy. Our results suggest that the DH31_CA_ neuron-CA axis modulates JH biosynthesis in response to dormancy-inducing conditions, revealing a regulatory neuroendocrine mechanism underlying reproductive dormancy.

Our study confirmed that DH31_CA_ neurons are identical to CA-LP1 and CA-LP2 neurons, which have been anatomically characterized in *D. melanogaster* larvae (44). Previous studies have indicated that CA-LP1 and CA-LP2 neurons have a negative effect on JH biosynthesis in male genital rotation during pupal development (45), consistent with the inhibitory role of DH31_CA_ neurons in JH biosynthesis during the adult stage. Nevertheless, we found that there was no phenotype of male genitalia rotation in genetic mutant males exhibiting either loss of *Dh31* or *Dh31-R* functions (Fig. S9A, B). These results indicate that, at least in the larval and/or pupal stages, DH31_CA_ neurons might produce an additional neuropeptide or neurotransmitter, different from DH31, which also has a negative impact on JH biosynthesis in the CA. Therefore, a complete list of the neuropeptides/neurotransmitters produced in DH31_CA_ neurons is required.

A recent study reported that DH31 and DH31-R were not involved in *D. melanogaster* oogenesis under non-dormancy-inducing conditions (71). Therefore, considering this report and the results of our study, it seems that DH31_CA_ neurons are essential only under dormancy-inducing conditions. However, whether and how DH31_CA_ neurons are regulated in response to dormancy-inducing conditions has not been elucidated. Previous studies have reported that the circadian clock system is involved in the regulation of reproductive dormancy in many insect species (4, 53, 72, 73), including *D. melanogaster* (27, 51, 60, 74). Therefore, we propose that, to an extent, circadian clock neurons, such as 5^th^ s-LNv, s-LNv, LNd, and DN1p are connected with DH31_CA_ neurons and modulated its activity under dormancy-inducing conditions. In addition, sNPF secreted from s-LNv may stimulate sNPF-R to modulate the activity of DH31_CA_ neurons.

Remarkably, a recent study showed that sNPF secreted from s-LNv cells positively regulates insulin-producing cells (IPCs), leading to the suppression of reproductive dormancy in *D. melanogaster* (51). Therefore, an alternative hypothesis is that sNPF from s-LNv simultaneously regulates DH31_CA_ neurons and IPCs to control reproductive dormancy. If sNPF plays a suppressive role in reproductive dormancy, it should negatively regulate DH31_CA_ neurons. Notably, the inhibitory effect of sNPF on central neurons has been reported in *D. melanogaster* (75).

This study, for the first time, identified DH31 as a neurotransmitter derived from neurons that directly innervate the CA. In contract, recent studies using *D. melanogaster* have revealed a number of circulating factors crucial for regulating reproductive dormancy (5). Most of these studies have repeatedly emphasized the importance of insulin-like peptides (ILPs) produced in IPCs located in PI. ILPs directly stimulate CA to enhance JH biosynthesis through the PI3-kinase-mTOR pathway under non-dormancy-inducing conditions (30). Conversely, under dormancy-inducing conditions, the production and secretion of ILPs from IPCs are inhibited, resulting in the suppression of JH biosynthesis, thus leading to reproductive dormancy (5). The modulation of IPC activity is mediated by multiple neuropeptides and neurotransmitters, including PDF and sNPF produced by the s-LNv cells, as well as monoaminergic neurotransmitters, such as serotonin and dopamine (29, 51). Dopamine also suppresses JH biosynthesis via a dopamine receptor subtype in CA (29). On the other hand, the release of allatostatin C (AstC) from DN3s is essential for cold-induced reproductive dormancy through the activation of cholinergic AstC receptor type 2 in clock neurons (76). DH31_CA_ neurons might cooperate with these humoral factors for the proper regulation of reproductive dormancy *in vivo*; however, it is not known how all these signals are integrated to regulate reproductive dormancy.

A previous study has reported that cAMP-dependent protein kinase, also known as protein kinase A (PKA), is required for JH biosynthesis (29). Considering this and our data from PinkFlamindo imaging, it is plausible that DH31 regulates JH biosynthesis by modulating intracellular cAMP levels; however, additional studies are required to elucidate the molecular basis of cAMP and PKA modulation of JH biosynthesis. Based on our immunostaining data, DH31 signaling did not seem to affect the protein levels of JH acid *O*-methyl transferase (JHAMT), a critical enzyme in the JH biosynthesis pathway (77, 78).

JH plays key roles in regulating numerous *D. melanogaster* adult female physiological processes, including vitellogenesis in non-dormancy-inducing conditions, ovulation, egg shape maintenance, gut remodeling, innate immunity, sleep regulation, and aging (64, 68, 79–83). In addition, JH signaling is also essential for regulating adult male physiological processes in *D. melanogaster*, including protein synthesis in accessory glands, sleep regulation, courtship motivation, and courtship-associated memory retention (82, 84–87). In fact, we found that DH31_CA_ neurons are also present in adult males (Fig. S9C) and therefore assume that DH31_CA_ neurons also regulate male adult JH biosynthesis. Further studies could reveal additional roles of DH31_CA_ neurons in modulating JH-dependent events in adult male and female flies.

Considering that DH31 is a well conserved molecule in invertebrates (88), future studies could also investigate the role of DH31_CA_ neurons in modulating JH-mediated reproductive dormancy in other insects. In particular, the morphology of *D. melanogaster* DH31_CA_ neurons closely resembled that of the CA-projecting PL neurons in *P. Terraenovae* (15). Therefore, DH31_CA_ neurons appear to be involved in the regulation of reproductive dormancy in other dipteran species.

Our study provides conclusive genetic evidence that *D. melanogaster* female reproductive dormancy is regulated by DH31-producing neurons that project to the CA. Further functional analysis of the clock neurons–DH31_CA_ neurons–CA axis in other insects could shed light on the conserved molecular mechanisms underlying the environmental adaptation of insects, including agricultural pests and infectious disease vectors. These studies would provide important information that will help establish the foundation for the development of new approaches for the control of harmful insects. Lastly, as DH31 belongs to an evolutionarily conserved peptide family that includes mammalian calcitonin gene-related peptides (89), it would be intriguing to examine whether these related peptides are also involved in regulating dormancy or hibernation in other animal species.

## Materials and Methods

### Drosophila culture

*Drosophila melanogaster* flies were raised until eclosion on a standard yeast-cornmeal-glucose fly medium (0.275 g agar, 5.0 g glucose, 4.5 g cornmeal, 2.0 g yeast extract, 150 μL propionic acid, and 175 μL 10% butyl p-hydroxybenzoate (in 70% ethanol) in 50 mL water) at 25 °C under a 12:12 h light/dark cycle. To induce reproductive dormancy, adult virgin females were collected within 6 h of eclosion and reared on standard medium at 11 ± 0.5 °C under short-day length (SD: 10:14 h light/dark cycle) conditions for 12 d, as described in previous studies (29, 30). To analyze the effects of JH analogs on reproductive dormancy, virgin female flies were collected within 6 h of eclosion and reared on standard medium supplemented with 1.5 mM methoprene (Sigma-Aldrich, St Louis, MO, PESTANAL 33375, racemic mixture; 1.5 M stock was prepared in ethanol) or 0.1% ethanol (control).

### Drosophila stocks

The following *Drosophila* genetic mutant strains were used: *Dh31*^*#51*^ (33) (from Fumika N. Hamada, Cincinnati Children’s Hospital Medical Center, USA), *Dh31*^*attp*^ (90) BDSC #84490), and *Dh31-R*^*attp*^ (90); BDSC #84491) (from Yi Rao, Peking University School of Life Sciences, China).

Additionally, the following transgenic *D. melanogaster* strains were used: *Aug21-GAL4* (44) (BDSC #30137), *Df(2R)BSC273* (BDSC #23169), *Dh31-R-T2A-GAL4* (91), *GMR21C09-GAL4* (BDSC #48936), *GMR94H10-GAL4* (BDSC #47268), *JHAMT-GAL4* (84) from Brigitte Dauwalder, University of Houston, USA), *JHMAT-LexA* (from Naoki Yamanaka, U.C. Riverside, USA), *Kurs21-GAL4* (44); from Stéphane Noselli, Université Côte D’Azur, France), *LexAop-CD4::spGFP11,UAS-CD4::spGFP1-10* (BDSC #58755), *Pdfr-T2A-GAL4* (91), *UAS-Dh31* (FlyORF #F003632), *UAS-Dh31-IR* (BDSC #41957), *UAS-Dh31-R-IR*^*NIG*^ (NIG-FLY #17043R-1), *UAS-Dh31-R-IR*^*KK*^ (VDRC #101995), *UAS-Dicer2* (BDSC #24651), *UAS-mCD::GFP* (BDSC #32219), *UAS-DenMark, UAS-Syt1::GFP* (BDSC #33064), *UAS-PinkFlamindo* (this study), *sNPF-R-T2A-GAL4* (BDSC #84691), *Viking-GFP* (DGRC #110692), and *UAS-GFP, mCD8::GFP* (92) from Kei Ito, University of Cologne, Germany), Heterozygous controls were obtained by crossing *w*^*1118*^ with strains of genetic mutants, GAL4 drivers, or UAS effectors.

### Generation of mouse anti-DH31 antibody

Peptides corresponding to the C-terminal 16 amino acid sequence (NH2-AKHLMGLAAANFAGGP-NH2) of *Bombyx mori* DH31 (GenBank accession No. BAG49567.1), with an N-terminal addition of cysteine, were synthesized and conjugated with maleimide-activated BSA (Imject Maleimide-Activated BSA, Thermo Fisher Scientific, 77115). The BSA-conjugated DH31 partial peptides were dialyzed with phosphate buffer saline (PBS). Subsequently, 50 μL of 1 mg/mL dialyzed conjugates was mixed with 25 μL of ABISCO-100 adjuvant (Isconova, Uppsala, Sweden) and 25 μL of PBS. The mixture was subcutaneously injected twice into the mice and whole blood was collected 12 d after the last immunization. Blood serum was heat-inactivated at 56 °C for 30 min, followed by the addition of an equal volume of saturated ammonium sulfate solution to precipitate the proteins. The precipitate was dissolved and dialyzed twice in PBS. We would like to emphasize that there was only one amino acid difference in the C-terminal 16 amino acid sequence of DH31 between *B. mori* and *D. melanogaster* (NH_2_-AKHRMGLAAANFAGGP-NH_2_; GenBank accession No. NP_523514.1); therefore, cross-reactivity was expected.

### Immunohistochemistry

The tissues were dissected in PBS and fixed in 4% paraformaldehyde in PBS for 30–60 min at 25–27 °C. The fixed samples were rinsed thrice in PBS, washed for 15 min with PBS containing 0.3% Triton X-100 (PBT), and treated with a blocking solution (2% bovine serum albumin in PBT; Sigma #A9647) for 1 h at 25–27 °C or overnight at 4 °C. The samples were incubated with a primary antibody in a blocking solution overnight at 4 °C. The primary antibodies used were as follows: chicken anti-GFP antibody (Sigma #G6539, 1:2,000), rabbit anti-RFP antibody (Medical & Biological Laboratories PM005, 1:2,000), guinea pig anti-JHAMT antibody (43) 1:1,000–2,000); rabbit anti-JHAMT antibody (78) 1:1,000), mouse anti-DH31 antibody (this study; 1:200), guinea pig anti-DH31(31) from Michael Nitabach, Yale University, USA; 1:500), and rabbit antibody against PDF of the cricket *Gryllus bimaculatus* (93) 1:2000). Notably, previous studies have confirmed that anti-cricket PDF antibodies cross-react with *D. melanogaster* PDF protein (94, 95). The samples were rinsed thrice with PBS and then washed for 15 min with PBT, followed by incubation with fluorophore (Alexa Fluor 488, 546, or 633)-conjugated secondary antibodies (Thermo Fisher Scientific; 1:200) in blocking solution for 2 h at RT or overnight at 4 °C. After another round of washing, all the samples were mounted on glass slides using FluorSave reagent (Merck Millipore, #345789).

### Connectome analysis

Connectome analysis was performed using NeuronBridge (96–98) https://neuronbridge.janelia.org/) and neuPrint+ (99, 100) https://neuprint.janelia.org/?dataset=hemibrain:v1.2&qt=findneurons). NeuronBridge is a database in which neurons labeled by GAL4 drivers are mapped after identification using light and electron microscopic connectome analysis. The NeuroBridge body IDs corresponding to two of the three pairs of DH31_CA_ neurons (Fig. 2A) were #295063181 and #5813067334. Neurons connected with #295063181 and #5813067334 were identified in the connectome database on neuPrint+; the names of the connected neurons and data of the connection numbers were obtained. The percentages of neurons indicate the proportions of each clock neuron in relation to the whole connection to #295063181 and #5813067334 (Fig. 2A).

### Counting mature egg numbers in ovaries

The ovaries of virgin females were dissected in PBS. The numbers of mature eggs (stage-14 oocytes) (101) in the ovaries were counted under a stereomicroscope (Leica MZ10F).

### *Generation of UAS-PinkFlamindo* transgenic line

The pcDNA3.1 plasmid containing the PinkFlamindo coding sequence (PinkFlamindo-pcDNA3.1; (59) was obtained from Addgene (#102356). The following primers were used for PCR amplification, to obtain the PinkFlamindo coding sequences with *EcoR*I and *Xba*l sites at the 5’ and 3’ termini, respectively; PinkFlamindo F (5’-ACTGAATTCATGCTGGTGAGCAAGGGC-3’) and PinkFlamindo R (5’-CCTGCTCGACATGTTCATTAGATCTCAG-3’). PCR products were digested with *EcoR*I and *Xba*I, purified, and ligated with *EcoR*I-*Xba*I the EcoRI-XbaI-digested pWALIUM10-moe plasmid (102). Transformants were generated using the phiC31 integrase system in the *P{CaryP}attP2* strain (103) by WellGenetics, Inc. w+ transformants of pWALIUM10-moe were established using standard protocols.

### cAMP imaging in the CA

In the implementation of live imaging in the CA, experimental conditions were optimized with reference to previous studies (85, 104). cAMP transients in the CA were imaged in flies expressing *UAS-PinkFlamindo* and *UAS-mCD8::GFP* driven by *JHAMT-GAL4*. Newly eclosed virgin females were cultured at 25 °C under a 12:12 h light/dark cycle for 1 d. Adult brain-gut complexes were dissected in Schneider’s Drosophila Medium (SDM; Thermo Fisher Scientific #21720024) without supplementation with fetal bovine serum. The brain-gut complexes were then treated with 100 μL of collagenase solution (0.05 mg/mL collagenase in SDM; Sigma-Aldrich #C0130) with gentle rotation for 9 min at 25–27 °C, and vortexed gently for 1 min ensuring that the brain and gut were not physically separated. The samples were then washed twice with 1 mL of SDM and once with 500 μL of SDM. The dissected tissues were held in a glass-bottom dish (35 × 10 mm, IWAKI #3910-035) with an insect pin (φ0.10 mm, Ento Sphinx Insect Pins) and silicone grease (Beckman), and 20 μL of SDM was added to cover the tissue. Live imaging was performed at 25–27 °C using a laser scanning confocal microscope (LSM700, Carl Zeiss) with a 20× objective lens. mCD8::GFP and PinkFlamindo were excited with 488 nm and 555 nm lasers, respectively. Time-lapse images were acquired every 8 s for 472 s. One-hundred and12 s after starting live-imaging, 80 μL of SDM with or without 375 nM synthetic DH31 peptide or 125 μM NKH477 was applied on the tissue. For image processing, CA was selected in a region of interest (ROI) over multiple time frames. The mean fluorescence intensities were measured along the time axis using the ImageJ software version 1.53q (105). Data were analyzed using Microsoft Excel.

### Measurement of juvenile hormone III titers

Newly emerged virgin females were collected within 6 h of eclosion and reared on standard medium at 11 ± 0.5 °C under short-day length (SD: 10 L:14D) conditions for 6 d. Forty adult female flies of each genotype were punctured using a tungsten needle and placed in a plastic tube with a hole at the bottom. The tube was then connected to a silanized glass vial (GL Science # 5183-4507) and centrifuged at 9,100×*g* for 5 min. Pre-cooled PBS (150 µL) was added to the glass vial where the hemolymph was collected, and 6.25 pg/µL (in acetonitrile) of JH III-D3 (Tronto Research Chemicals #E589402) was added as an internal control. Subsequently, 600 µL of hexane was added, and the samples were stirred for 1 min. The samples were then centrifuged at 2,000×*g* for 5 min at 4 °C, and 500 µL of the organic phase was transferred to a fresh silanized vial. The samples were dried under a gentle nitrogen flow and stored at -20 °C until further analysis. JH titers from these hemolymph extracts were determined using liquid chromatography coupled with electrospray tandem mass spectrometry (LC-MS/MS) as previously described (63).

### Reverse transcription-quantitative PCR (RT-qPCR) for yolk protein expression

Total RNA was extracted from whole bodies of 6-day-old adult virgin female flies (under a 10:14 h light/dark cycle) after approximately 4–6 h of light period. RNA was reverse-transcribed using ReverTra Ace qPCR RT Master Mix with gDNA Remover (TOYOBO). Synthesized cDNA samples were used as templates for quantitative PCR using THUNDERBIRD SYBR qPCR Mix (TOYOBO) on a Thermal Cycler Dice Real Time System (Takara Bio). The amount of target RNA was normalized to the endogenous control *ribosomal protein 49* gene (*rp49*) and the relative fold change was calculated. The expression levels of *yolk protein 1, yolk protein 2*, and *yolk protein 3* were compared using the ΔΔCt method. The following primers were used for this analysis: rp49 F (5’-CGGATCGATATGCTAAGCTGT-3’), rp49 R (5’-GCGCTTGTTCGATCCGTA-3’), yolk protein1 F (5′-CAGGCTCAGTACACCCACAC-3′), yolk protein1 R (5′-CTCAACGTTGTGGTGGATCTG-3′), yolk protein2 F (5’-ACCCTTAAGAAGCTGCAGGAG-3’), yolk protein2 R (5’-ATGGTTGAACTGGGACAGATG-3’), yolk protein3 F (5’-CTCAAGAGCAGCGACTACGAC-3’), and yolk protein3 R (5’-TAGCGTTTGAAGTTGGTCAGG-3’).

### Statistical analysis

The sample sizes were chosen based on the number of independent experiments required for statistical significance and technical feasibility. The experiments were not randomized, and the investigators were not blinded. All statistical analyses were performed using the “R” software version 4.0.3. Details of the statistical analyses are described in figure legends.

## Acknowledgments

We thank Brigitte Dauwalder, Fumika N. Hamada, Kei Ito, Stéphane Noselli, Yi Rao, Naoki Yamanaka, Bloomington Drosophila Stock Center, Vienna Drosophila Resource Center, and National Institute of Genetics (NIG-FLY) for fly stocks; Outa Uryu, Michael Nitabach, Yoshiaki Tanaka, and Kenji Tomioka for antibodies; Addgene for plasmids; WellGenetics for generating transgenic flies; and Editage (www.editage.com) for English language editing. We are also grateful to Kazuki Harada and Takashi Tsuboi for their advice on the PinkFlamindo experiment; Chen Zhang and Young-Joon Kim for their advice on JH titer measurement; Tomotsune Ameku and Yuto Yoshinari for their advice on the fly works; and Masaharu Hasebe and Sakiko Shiga for critical reading of the manuscript. Y.K., E.I., and R.H. received a fellowship from the Japan Society for the Promotion of Science (JSPS). This work was supported by JSPS KAKENHI (21J20365 to Y.K., 17J00218 to E.I., 26250001 and 17H01378 to H.T.) and a Grant-in-Aid for Scientific Research on Innovative Areas (21H00226) from JSPS to R.N. This research was also funded by project 22-21244S from the Czech Science Foundation, Czech Republic, to M.N., and grants R21AI167849 to F.G.N. and R21AI153689 to M.N. from the National Institutes of Health-NIAID, USA.

## Notes

### Competing Interest Statement

The authors have declared no competing interest.

