## Supplemental Figures S1-S9 for "Female reproductive dormancy in *Drosophila melanogaster* is regulated by DH31-producing neurons projecting into the *corpus allatum*"

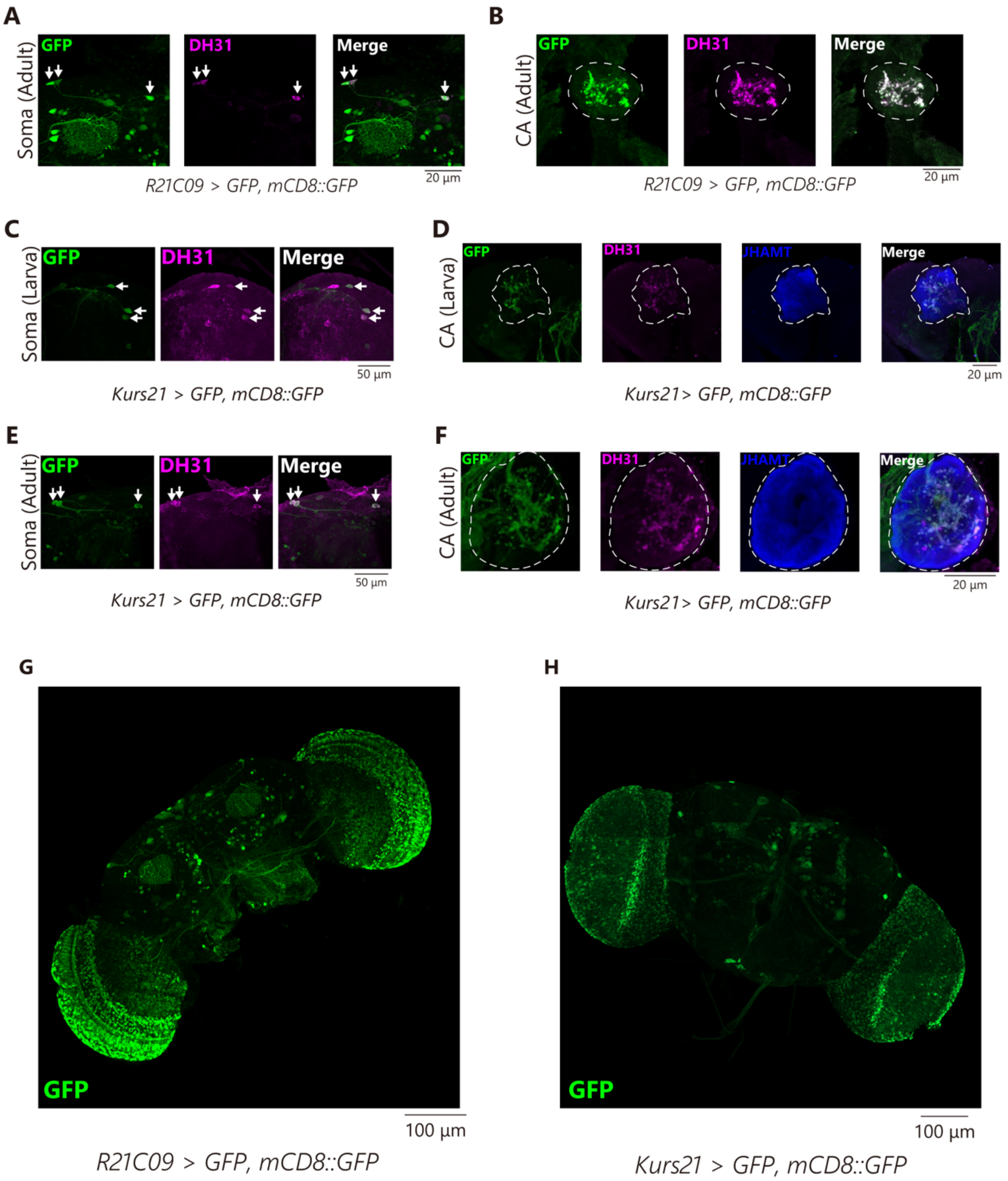


**Figure S1. *R21C09-GAL4* and *Kurs21-GAL4* are active in DH31$_{CA}$ neurons.**

(A, B) GFP signals in the soma (A) and CA region (B) of an adult female *R21C09-GAL4 UAS-GFP UAS-mCD8::GFP*. Samples were immunostained with anti-GFP (green) and anti-DH31 (magenta) antibodies. Arrows indicate the soma of DH31$_{CA}$ neurons and dashed lines outline the CA.

(C, D) GFP signals in the soma (C) and CA region (D) of a larva of *Kurs21-GAL4 UAS-GFP UAS-mCD8::GFP*. (C) Immunostaining with anti-GFP (green) and anti-DH31 (magenta) antibodies. Arrows indicate the soma of $DH31_{CA}$ neurons. (D) Immunostaining with anti-GFP (green), anti-DH31 (magenta), and anti-JHAMT (blue) antibodies.
(E, F) GFP signals in the soma (E) and CA regions (F) of an adult female *Kurs21-GAL4 UAS-GFP UAS-mCD8::GFP*. (E) Immunostaining with anti-GFP (green) and anti-DH31 (magenta) antibodies. Arrows indicate the soma of $DH31_{CA}$ neurons. (F) Immunostaining with anti-GFP (green), anti-DH31 (magenta), and anti-JHAMT (blue) antibodies.
(G, H) Lower magnification views of *R21C09-GAL4 UAS-GFP UAS-mCD8::GFP* (G) and *Kurs21-GAL4 UAS-GFP UAS-mCD8::GFP* (H). Samples were immunostained with an anti-GFP antibody (green).

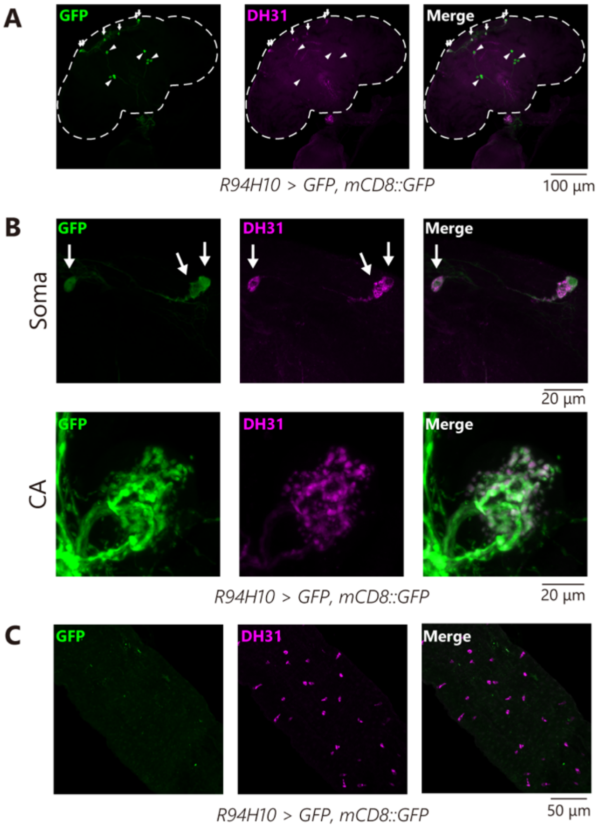


**Figure S2. Expression pattern of *R94H10-GAL4*.**

(A) Transgenic visualization of $DH31_{CA}$ neurons by GFP driven by *R94H10-GAL4*. The sample is the same as that shown in Fig. 1D, while signals were visualized with anti-GFP (green) and anti-DH31 (magenta) antibodies. Note that some *R94H10-GAL4*-positive neurons marked with arrowheads, which do not innervate the CA, were DH31-negative.

(B) Immunostaining signal with anti-GFP (green) and anti-DH31 (magenta) antibodies in the brain region, including the soma of $DH31_{CA}$ neurons (arrows), and in the CA region in an adult female of *R94H10-GAL4 UAS-GFP UAS-mCD8::GFP*.

(C) Immunostaining signal with anti-GFP (green) and anti-DH31 (magenta) antibodies in the posterior midgut of an adult female of *R94H10-GAL4 UAS-GFP UAS-mCD8::GFP*.

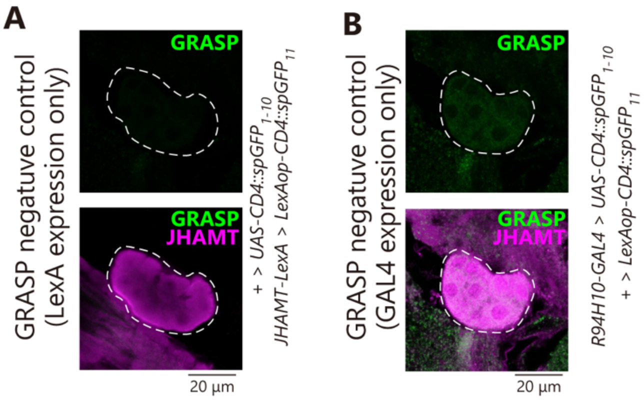


**Figure S3. GFP signals of GRASP negative control adult females.**

See Fig. 1F for GRASP positive signals.

(A) GRASP signals in the CA region of a virgin female expressing *LexA* in the CA but not *GAL4* in DH31$_{CA}$ neurons (+/*JHAMT-LexA; +/UAS-CD4::spGFP*$_{1-10}$ *LexAOP>CD4::spGFP*$_{10}$).

(B) GRASP signals in the CA region of a virgin female expressing *GAL4* in DH31$_{CA}$ neurons but not *LexA* in the CA (+/+*; R94H10-GAL4/UAS-CD4::spGFP*$_{1-10}$ *LexAOP>CD4::spGFP*$_{10}$).

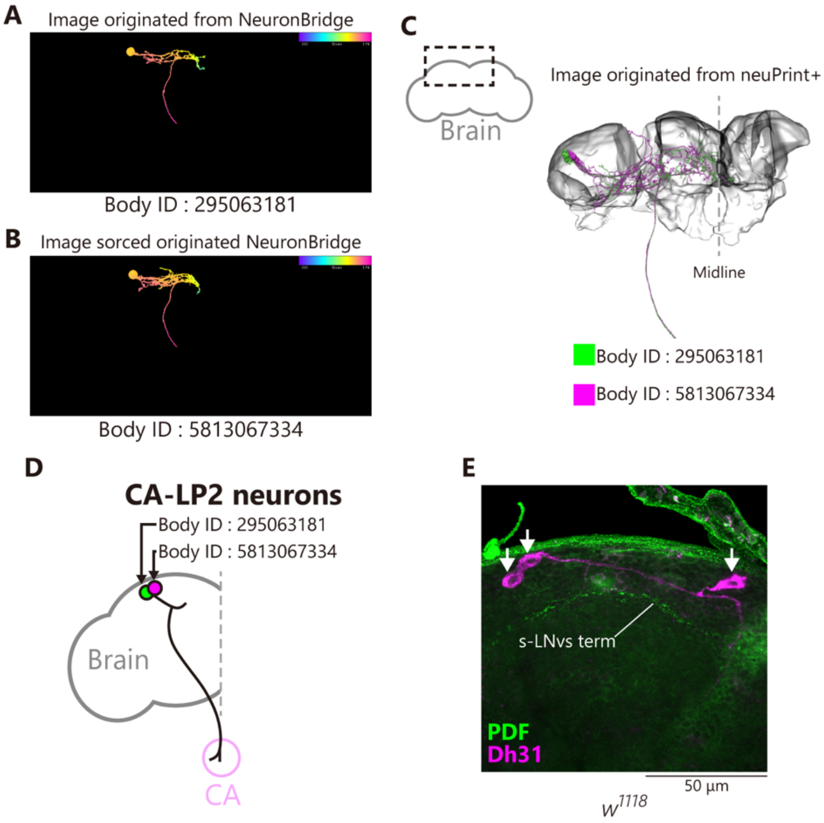


**Figure S4. Computational identification of CA-LP2 neurons.**

Supplementary data for Fig. 2A.

(A, B) Computationally-reconstructed images of the two pairs of $DH31_{CA}$ neurons. There images were generated using NeuronBridge (96–98) and deposited in the publically-available website (https://neuronbridge.janelia.org/). We used the reconstructed image of Body IDs 29506318 (A) and 5813067334 (B) under the condition of Creative Commons License CC-BY 4.0. Relative positions of cell bodies and neurites are represented in color bar scales, in which a surface or a deep area of the brain are represented with blue and red, respectively.

(C) A computationally-merged reconstructed image of Body IDs 29506318 (green) and 5813067334 (magenta). The image on the right side, generated using neuPrint+ (99, 100), was derived from the publically-available website (https://neuprint.janelia.org/?dataset=hemibrain:v1.2&qt=findneurons). This neuPrint+ image corresponds to the dorsolateral and dorsomedial brain region, marked by dashed lines in the whole brain illustration (left side). We use the image under the condition of Creative Commons License CC-BY 4.0.

(D) A schematic representation of the relative positions of Body IDs 29506318 (green) and 5813067334 (magenta) in the brain hemisphere based on C. Considering the positions of the cell bodies, we concluded that the neurons of Body IDs 29506318 and 5813067334 correspond to CA-LP2 neurons.

(E) Immunostaining with anti-PDF (green) and anti-DH31 (magenta) antibodies in the brain region, including the soma of $DH31_{CA}$ neurons (arrows), of wild-type ($w^{1118}$) adult females. Axonal termini of s-LNv neurons (s-LNv term) are indicated by ‘s-LNvs term.’

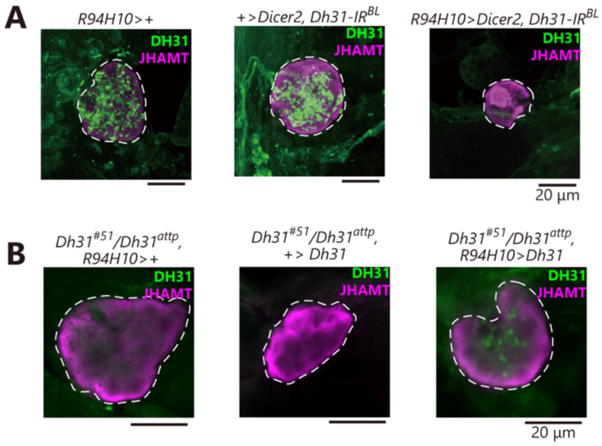


**Figure S5. DH31 immunoreactive signals in the CA region of several genotypes.**

Supplementary data for Fig. 3.

Samples were derived from virgin females 12 d after transfer to dormancy-inducing conditions. (A) Immunostaining signals of the anti-DH31 antibody (green) in the CA region of control and $DH31_{CA}$ neuron-specific *Dh31* RNAi adult females. CA was visualized using an anti-JHAMT antibody (magenta). DH31 puncta were diminished in the CA region of *Dh31* RNAi animals. (B) Immunostaining signals of the anti-DH31 antibody (green) in the CA region of *Dh31* transheterozygous mutant females in the presence or absence of $DH31_{CA}$ neuron-specific expression of *Dh31* cDNA under dormancy-inducing conditions. DH31 puncta were recovered in the CA region of the transgenic rescue animals.

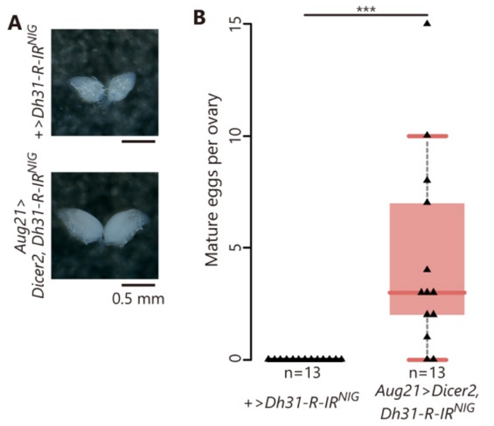


**Figure S6. RNAi of DH31 receptor in the CA leads to failure in the induction of reproductive dormancy.**

Supplementary data for Fig. 4. Samples were derived from virgin females 12 d after the transfer to dormancy-inducing conditions.

(A, B) Mature egg formation in the control and CA-specific *Dh31-R* RNAi females under dormancy-inducing conditions. These data were obtained from the transgenic RNAI strain from the National Institute of Genetics (NIG). (A) Representative images of the ovaries. (B) Quantification of mature eggs per ovary.

Statistical analysis: Wilcoxon rank-sum test for B. ****P* <0.001. n.s.: not significant.

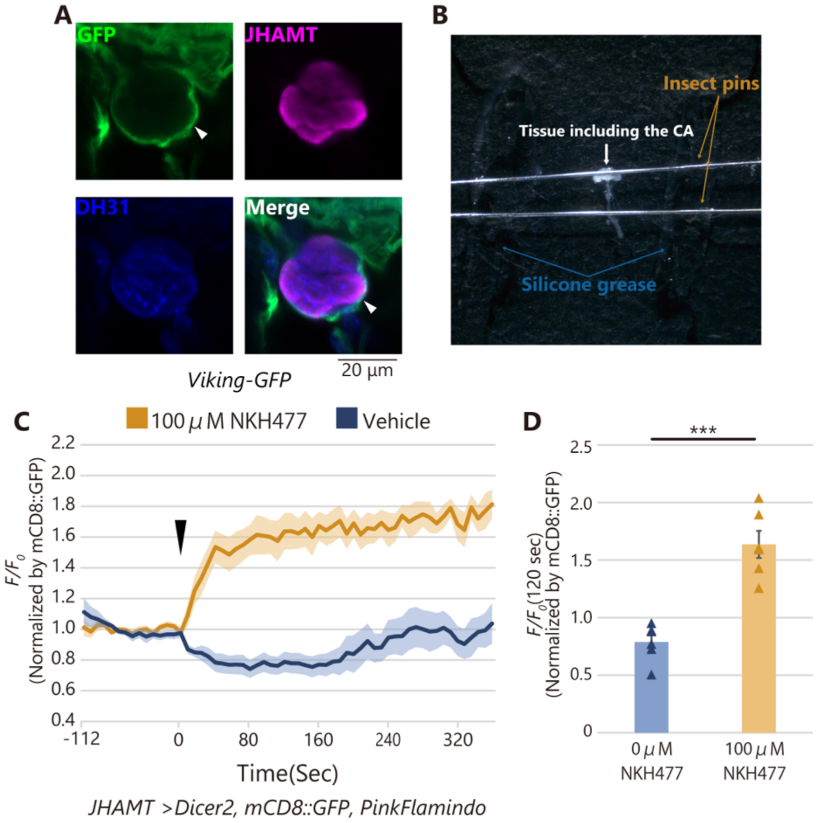


**Figure S7. PinkFlamindo imaging in the CA.**

Supplementary data for Fig. 5.

(A) A representative photograph demonstrating how to mount the dissected, unfixed tissue containing the CA. To keep the tissue immobile, two insect pins were placed on the tissue, being careful not to hide the CA. To fix the position of the insect pins, silicon grease is applied to the bottom of the dish.

(B) GFP signal (green) in the CA of *Viking-GFP* strain that produces a functional GFP-tagged version of Viking protein. The sample was immunostained with anti-JHAMT (magenta) and anti-DH31 (blue) antibodies. The arrowhead indicates the presence of collagen layer surrounding the CA.

(C) Changes in the relative fluorescence intensity of *JHAMT-GAL4*-driven PinkFlamindo in the CA after 360 s with or without stimulation by 100 μM NKH477 (each n = 6). $F/F_0$ values are normalized by signal intensity of *JHAMT-GAL4*-driven mCD8::GFP.

(D) Quantification of normalized PinkFlamindo signals in the CA at 120 s after the stimulation. Values in C and D are presented as mean ± SE. Statistical analysis: Student's *t*-test for D. ****P* <0.001. n.s.: not significant.

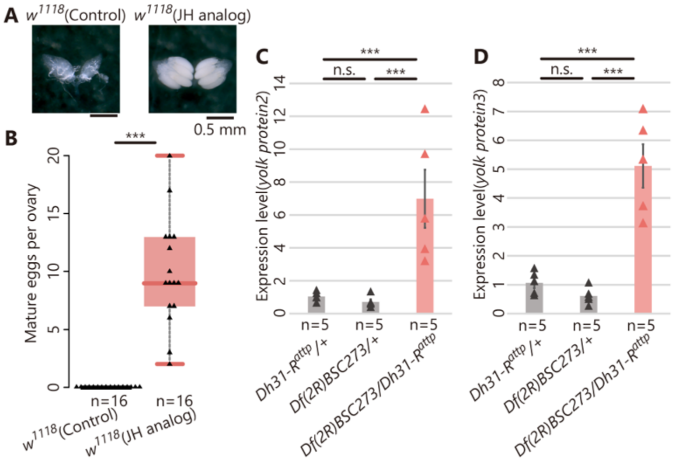


**Figure S8. The effect of JH analog on mature egg formation and mRNA levels of *yolk protein 2* and *yolk protein 3* under dormancy-inducing conditions.**

Samples were derived from virgin females 6 d (for A and B) or 12 d (for C and D) after transfer to dormancy-inducing conditions.
(A, B) Mature egg formation in wild-type ($w^{1118}$) adult females, with or without oral administration of methoprene (JH analog, JHA) under dormancy-inducing conditions. (A) Representative images of the ovaries. (B) Quantification of mature eggs per ovary.
(C, D) Quantification of mRNA of *yolk protein 2* (C) and *yolk protein 3* (D) in control and *Dh31-R* genetic mutant females under dormancy-inducing conditions by RT-qPCR.
Values in C and D are presented as mean ± SE. Statistical analysis: Wilcoxon rank sum test for B; Tukey–Kramer's HSD test for C and D. ***$P$ <0.001. n.s.: not significant.

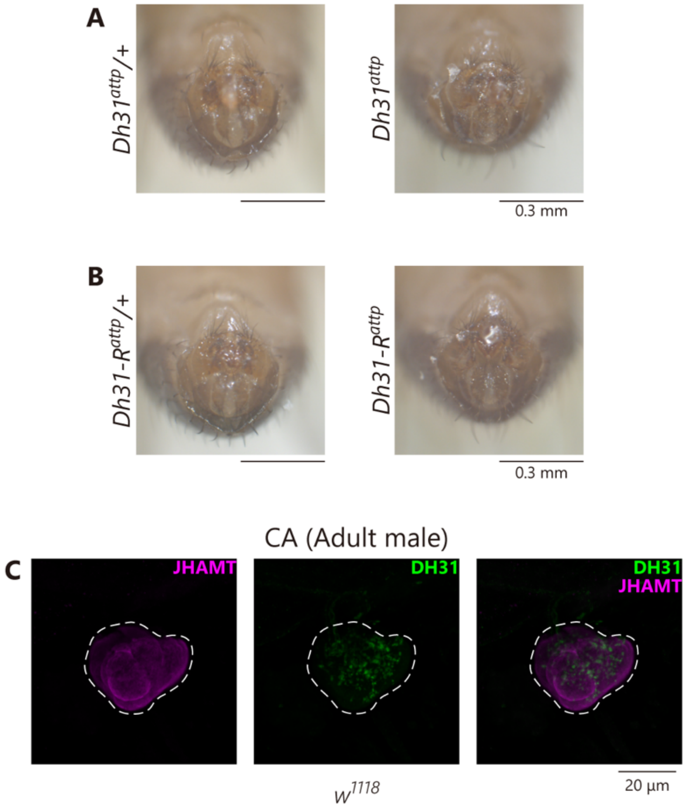


**Figure S9. Male genitalia of loss of *Dh31* and *Dh31-R* genetic mutants, and DH31$_{CA}$ neurons in the adult male.**

(A, B) Male genitalia of loss of *Dh31* (A, right) and *Dh31-R* (B, right) genetic mutants. Control heterozygous animals are shown in the left side.

(C) Immunostaining signal with anti-DH31 antibody (green) along with anti-JHAMT antibody to visualize the CA (magenta) of wild-type ($w^{1118}$) adult male. Note that DH31-positve puncta are observed in the CA region.
